# Measurement of nitric oxide production in mouse tracheal epithelial cell cultures differentiated at the air liquid interface

**DOI:** 10.1101/2025.11.23.690029

**Authors:** Muddassar Iqbal, David Mahan, Adam Clark, Akueba Bruce, Anthony W. DeMartino, Paola Corti

**Affiliations:** Department of Medicine, University of Maryland School of Medicine, Baltimore MD 21201; Department of Biochemistry and Molecular Biology, University of Maryland School of Medicine, Baltimore MD 21201

**Keywords:** Nitric Oxide (NO), Airway cilia, Mouse tracheal epithelial cells (MTEC), Air-liquid interface (ALI), Apical mucus, Nitric Oxide Analyzer (NOA)

## Abstract

Airway ciliary dysfunction often leads to impaired mucociliary clearance, a defect frequently associated with dysregulation of canonical nitric oxide (NO) signaling. Indeed, many airway diseases exhibit aberrant levels of exhaled NO, a feature commonly used as a diagnostic indicator in patients. While this measurement is routinely performed clinically, there is currently no validated approach to assess NO release from airway ciliated cells in vitro, limiting mechanistic studies of NO regulation and function. An established method to study airway diseases in vitro is the use of cultured mouse tracheal epithelial cells (MTECs) differentiated at the air–liquid interface (ALI). Here, using this model, we develop a protocol to measure NO released from ciliated cells and accumulated in apical mucus, collected from the ALI-differentiated cultures. Using established instrumentation for NO analysis and a triiodide-based assay, we quantify nitrite, a stable NO metabolite and biomarker of NO production. For assay validation, cells were treated with the NO donor Diethylenetriamine NONOate (DETA-NONOate or DETA/NO) or the NO synthase inhibitor N(ω)-Nitro-L-arginine methyl ester (L-NAME). Together, these approaches demonstrate that this method reliably detects NO in samples from ALI-differentiated airway epithelia, providing an accurate in vitro platform to quantify NO release and model diseases associated with abnormal NO metabolism.

**SUMMARY:** The goal of this current protocol is to describe a cell culture-based method to study airway diseases associated with altered nitric oxide (NO) signaling. This method describes the quantification of NO in airway epithelia differentiated at the air-liquid interface (ALI) using a triiodide-based chemiluminescence assay.

## INTRODUCTION

A hallmark of several airway diseases, including asthma, chronic obstructive pulmonary disorder (COPD), cystic fibrosis (CF) and primary ciliary dyskinesia (PCD), is altered nitric oxide (NO) homeostasis measurable as changes in exhaled NO levels, which can largely vary in these diseases, being elevated in COPD and asthma^**1-2**^, while drastically reduced in CF and PCD^**3-4**^. These diseases commonly exhibit abnormalities in airway epithelial motile cilia, the key effectors of mucociliary clearance^**5**^. Because NO and its downstream signaling pathways are critical regulators of ciliary motility, understanding how NO production and signaling are altered in these contexts is essential for elucidating mechanisms of mucociliary dysfunction. Accordingly, a method that enables precise detection of NO production and release in airway epithelia provides a valuable tool for dissecting the regulation and physiological role of this signaling network.

NO regulates airway cilia biogenesis and function by influencing ciliated cell polarity and the coordinated beating of motile cilia^**6-8**^. However, the mechanisms that maintain NO homeostasis in the airways remain incompletely understood. Airway epithelial cultures differentiated at air– liquid interface (ALI) provide a valuable in vitro system for investigating these mechanisms. At present, this model is widely used to study the impact of environmental insults on the airway epithelium^**9-11**^ and to elucidate the pathophysiology of airway diseases^**12**^. Compared with conventional submerged cultures^**13-14**^, ALI-differentiated epithelia more faithfully reproduce the architecture and function of the native airway, including epithelial polarization, mucociliary differentiation, and realistic gas–liquid exchange, making them especially suitable to study apical NO release, signaling, and related ciliary function.

Accurate measurement of NO concentration is essential, as both excessive and deficient NO levels can impair ciliary function. Currently, fluorescent probe-based methods (e.g., diaminofluoresceins, DAFs) available for detecting NO are only semi-quantitative and are limited by pH sensitivity, oxygen dependence and lack of specificity, as they can react with cellular oxidants and antioxidants^**15**^. To overcome these limitations, we established a protocol to quantify NO in apical secretions collected from ALI-differentiated airway epithelia using a Nitric Oxide Analyzer (NOA). The NOA is a highly sensitive chemiluminescence-based detector traditionally used to measure NO metabolites in chemical samples and more recently adapted for the analysis of biological specimens, including blood, tissue homogenates, and cell culture supernatants^**7,16-17**^. NO is a highly reactive signaling molecule rapidly oxidized to nitrite in tissues under conditions of normal oxygen levels, with an exceptionally short half-life. Thus, nitrite as end product of NO oxidation serves as an indicator of total NO production in biological systems^**18-20**^. This sensitive and reproducible chemiluminescence method allows detection of nitrite/NO at picomolar levels^**21-22**^, making it ideal for measuring small changes in low-volume biological samples such as those derived from MTEC ALI cultures.

## PROTOCOL

The protocol takes 2 days for isolation of mice tracheal epithelial cells (MTEC), 7-10 days for expansion, 5-6 days for proliferation in transwell inserts, 21 days for differentiation (time needed to acquire sufficient multi-ciliated cells), 24 hours of drug treatment and a day for sample collection, NOA calibration, injections and data acquisition.

### 1. Preparation of media and solutions

Note: See Table 1 for details on media ingredients.

**Table 1.** Media compositions for mouse tracheal epithelial cell (MTEC) culture and ALI differentiation. The media ingredients for MTEC expansion in T75 cell culture flask, proliferation in 12 mm (0.4 µm pore size) transwell inserts and differentiation at air liquid interface (ALI) are listed along with the vendor-provided or self-prepared stock and final concentrations.

|  | Reagent (Stock concentration) | Final concentration |
| --- | --- | --- |
| **MTEC Expansion media** | Keratinocyte serum-free media (1X) | 1X |
|  | Penicillin/Streptomycin (100X) | 1X |
|  | Murine epidermal growth factor (0.5 mg/ml) | 0.025 µg/ml |
|  | Bovine Pituitary Extract (15.1 mg/ml) | 0.03 mg/ml |
|  | Insulin-Transferrin-Selenium-Glutamine (100X) | 1X |
|  | (Add fresh) ROCK inhibitor, Y27632 (1 mM) | 10 µM |
|  | (Add fresh) Notch inhibitor DAPT (10 mM) | 5 µM |
| **MTEC Proliferation media** | DMEM/F12 with glutamine and HEPES (1X) | 1X |
|  | Fetal Bovine Serum | 2.50% |
|  | Penicillin/Streptomycin (100X) | 1X |
|  | Amphoterecin B (1000X) | 1X |
|  | Bovine pancreatic insulin 2 mg/ml) | 10 µg/ml |
|  | Murine epidermal growth factor (0.5 mg/ml) | 0.025 µg/ml |
|  | Bovine Pituitary Extract (15.1 mg/ml) | 0.03 mg/ml |
|  | Cholera toxin (0.1 mg/ml) | 0.1 µ/ml |
|  | Transferrin (5mg/ml) | 5 µg/ml |
|  | (Add fresh) ROCK inhibitor, Y27632 (1 mM) | 10 µM |
|  | (Add fresh) Retinoic acid (5 µM) | 0.05 µM |
| **MTEC Differentiation media** | DMEM/F12 with glutamine and HEPES (1X) | 1X |
|  | NuSerum | 2% |
|  | Penicillin/Streptomycin (100X) | 1X |
|  | Amphoterecin B (1000X) | 1X |
|  | (Add fresh) Retinoic acid (5 µM) | 0.05 µM |

1. Ham’s F-12 media Mix 500 ml F12 media (Ham) with 5 ml 100X Penicillin/Streptomycin antibiotics and 0.5 ml Amphoterecin B (Fungizone). Filter sterilize and store at 4°C.
2. Pronase solution Add 15 mg pronase to 10 ml Ham’s F12 media to make 0.15% pronase solution fresh before use.
3. DNase I solution Add 5 mg crude pancreatic DNase I to 9 ml Ham’s F12 media and 1 ml 10 mg/ml Bovine Serum Albumin (BSA) solution to prepare a 0.5 mg/ml DNase I solution. Make 1 ml aliquots and store at −20°C.
4. Ham’s F12 media with fetal bovine serum (FBS) Add 10 ml fetal bovine serum (FBS) to 40 ml Ham’s F12 media with antibiotics to make F12 media containing 20% FBS. Filter sterilize and store at 4°C.
5. MTEC Expansion media Prepare the MTEC expansion media (Table 1) and filter sterilize. Add ROCK inhibitor Y-27632 (5 mM stock solution) at 10 μM and Notch γ-secretase inhibitor DAPT (10 mM stock solution) at 5 μM fresh before use.
6. MTEC Proliferation media Prepare the MTEC proliferation media (Table 1) and filter sterilize. Before use add fresh Y-27632 (5 mM stock solution) at 10 μM and retinoic acid (5 μM stock) at 0.05 μM.
7. MTEC Differentiation media Prepare the MTEC differentiation media (Table 1) and filter sterilize. Add fresh retinoic acid at 0.05 μM before use.

### 2. Isolation, expansion and differentiation of mouse tracheal epithelial cells (MTEC)

1. Euthanize three adult mice (8-20 weeks old) according to the procedures following guidelines and regulations of the institution’s approved protocol.
2. Place the animal dorsally (chest facing up) on an aseptic surface. Using clean forceps, gently lift the skin and cut it open with surgical scissors from the lower lip to the abdomen.
3. Move the salivary glands to either side to expose the trachea. Remove the connective tissue and trachealis muscle to expose the cartilage rings.
4. Resect the entire length of trachea and place it in a Petri dish containing sterile Ham’s F12 medium.
5. Take the dish inside a laminar flow hood and transfer the tracheas to another dish with fresh Ham’s F12 medium.
6. Remove any extratracheal tissue and cut the trachea longitudinally through the lumen.
7. Place all the tracheas in a 50 ml screw-cap conical tube containing sterile 10 ml 0.15% pronase solution and incubate overnight at 4°C.
8. Coat a T75 cell culture flask with 10 ml collagen I (rat tail) at 50 μg/ml and incubate overnight at 37°C.
9. The next day invert the tube containing the tissues 10-15 times to mix and place back at 4°C for 45 min.
10. Stop the pronase reaction with 10 ml Ham’s media containing 20% FBS and invert the tube 10-15 times.
11. Add 10 ml Ham’s media with FBS in three 15 ml screw-cap conical tubes. With a Pasteur pipette transfer the tracheal tissue from the 50 ml tube to the first 15 ml tube and invert 10-15 times to dislodge the tracheal epithelial cells. Set the 50 ml tube aside on ice, repeat this transfer step with the other two 15 ml tubes and discard the tissue.
12. Combine the media from the three conical tubes with that in the 50 ml tube and centrifuge at 1400 RPM for 10 min at 4°C.
13. Carefully remove the supernatant and add 0.5 ml of 0.5 mg/ml DNase I solution. Incubate on ice for 5 min and then centrifuge at 1400 RPM for 5 min at 4°C.
14. Carefully discard the supernatant and resuspend the cell pellet in 7-10 ml MTEC expansion media.
15. Aspirate the collagen solution from the T75 flask and wash the flask three times with PBS (1X). Allow the flask to dry completely.
16. Seed the MTEC suspension into the flask and place in an incubator maintained at 37°C and 5% CO2.
17. Replace the expansion medium every 2 or 3 days and monitor increase in cell number.
18. Upon 70% confluency, detach the monolayer by adding 1 ml Accutase solution (Gibco), incubate for 10-15 min at 37°C.
19. Resuspend the cells in 10 ml of MTEC proliferation medium and spin at room temperature for 5 min.
20. Repeat the wash step, count cells, resuspend in proliferation media and seed 1-2 x 10^5^ cells in 0.5 ml volume per 12 mm transwell insert (0.4 μm membrane pore size) precoated with 50 µg/ml collagen overnight at 37°C.
21. Add 1.5 ml media in the basal chamber of each insert and place in the 37°C incubator.
22. Change media in both the apical and basal chamber on the third day and allow the cells to reach 100% confluency usually by day 5 or 6 of proliferation.
23. Once confluent, wash the cells apically with 0.5 ml MTEC differentiation media. To start ALI add 0.75 ml differentiation media only in the basal chamber leaving the cells in apical chamber dry.
24. Wash cells in the apical chamber with differentiation media every 2-3 days, replace media in the basal chamber and allow cells to differentiate at ALI for 21 days.

### 3. Immunolabeling and imaging of ciliated epithelial cells

1. At ALI day 7, 14 and 21, fix the cells with 4% paraformaldehyde (PFA) and immunolabel with anti-FoxJ1 antibody to detect FoxJ1, a transcription factor essential for ciliogenesis^**23-24**^, and anti-acetylated α-tubulin antibody to detect acetylated α-tubulin, a structural marker of motile cilia.
2. Image the immunolabelled cells through confocal microscopy (Nikon W1 spinning disk Ti2 inverted microscope with Hamamatsu sCMOS camera) and quantify the FoxJ1-positive or acetylated tubulin-positive cells as percentage of total (DAPI-positive) cells per region of interest (ROI) using the cell counter tool in FIJI imaging software.
3. Using the GraphPad Prism software (v. 10.0) prepare graphs and perform statistical analysis using Student’s t test (one-tailed).

### 4. Drug Treatment of differentiated airway epithelia

1. At ALI day 21 prepare 250 μM Diethylene triamine-NONOate (DETA/NO) in differentiation media by adding 7.5 μl DETA-NONOate from the 100 mM stock solution to media for a total volume of 3 ml. Add 750 μl of the final solution in three basal chambers of differentiated cells and incubate for 24 h at 37°C.
2. Prepare 1 mM Nω-Nitro-L-Arginine Methyl Ester (L-NAME) by adding 30 μl from the 100 mM L-NAME stock solution MTEC differentiation media for a total volume of 3 ml. Add 750 μl of the final solution in three basal chambers and incubate for 24 h at 37°C.
3. Add DMSO only (vehicle) in the basal media of three separate inserts to be used as control. For NOA analysis it is necessary to pull together samples from three inserts.

### 5. Sample collection

1. At ALI day 22, collect samples from the differentiating airway epithelia beginning with 750 μl of basal media from each of three inserts per experimental group.
2. Store immediately at −20°C until injection into NOA. Note: It is essential to also collect basal media incubated in inserts without cells, which serves as blank control when injected into the NOA to account for potential external nitrite contamination.
3. To collect the nitrite/NO present in the apical mucus of the epithelium, add 50 μl pre-warmed PBS (1X) buffer (sterile, aliquoted and frozen) on top of cells in each insert, incubate for 30 min at 37°C, then collect the PBS containing the mucus overlying the epithelium.
4. Combine the PBS washes from three inserts to obtain around 150 ul apical mucus/PBS from each experimental condition which will be enough for two NOA injections.
5. Store the apical wash samples at −20°C until further use. Note: As with the basal media samples, it is essential to collect and store the PBS wash buffer as the experimental samples for use as a blank control during NOA injections.
6. Wash the apical and basal sides of the epithelium with 500 μl prewarmed PBS twice.
7. Add 500 μl Accutase solution apically and incubate for 15-20 min until the cells completely detach from the membranes.
8. Once fully detached add serum-free RPMI-1640 media (at 3:1 media to Accutase ratio) and spin at 1400 RPM for 5 min at 4°C.
9. Resuspend the cell pellet in 50 μl lysis solution (RIPA buffer + Halt 1X protease inhibitor cocktail) and incubate on ice for 45 min.
10. Spin the lysate at 12,000 RPM for 20 min at 4°C and collect supernatant. Use 3 μl assay to determine the protein concentration through the BCA assay. Dilute the remaining lysate solution 1:1 in RIPA buffer to make enough volume for two 50 μl NOA injections.
11. Store the RIPA buffer at −20°C and the lysates at −80°C until injecting into the NOA.

### 6. NOA calibration and sample injections

1. Prepare triiodide reagent (400 mg potassium iodide, 250 mg iodine, 28 ml glacial acetic acid, 8 ml ultrapure water). Store at room temperature and use within two weeks of preparation.
2. Prepare 1 mM stock solution of sodium nitrite in nitrite-free water and freeze it at −20°C until further use. Dilute it fresh every time to prepare 0.125, 0.25, 0.5, 1.0, 2.0 and 4.0 μM nitrite standard solutions in PBS before the injections. Note: The PBS used here must be sterile and stored capped to avoid any nitrite contamination.
3. Turn on the nitrogen gas tank, increase the gas pressure very slightly and keep it low, usually close to 0 psi.
4. Assemble the glass Purge vessel and add 4 ml of triiodide reagent into the vessel with 20 μl anti-foam diluted 1:10. Securely cap it with a septum held in place by the screw cap to ensure an airtight seal.
5. Pour 10 ml sodium hydroxide (NaOH) into the NaOH trap.
6. On the display screen of the NOA machine (nCLD 88) select “Measurement”, “Select Gas and Range”.
7. Under Range (ppb) select “MR2:50.00” (highly sensitive setting used for very low nitrite in biological samples) and under Filter select “Fast”.
8. Adjust the gas pressure circulating in the apparatus until the reactor pressure on the NOA screen reads between 27-29.
9. On the computer screen coupled to the NOA open the eDAQ chart software.
10. On the upper right corner select the collecting speed as 10/s and the voltage to 10 V.
11. From the upper left tabs, click “Setup>Channel Settings”, select “Channel 1” and change the number at the bottom left to 1, click “OK”.
12. Set the scale on bottom right to 2:1. Press the start icon on the NOA instrument screen and click “Start” on the bottom right corner of the software.
13. Monitor the baseline. Once the baseline is stable, inject 10 μl of PBS, to ensure the baseline stays stable, through the septum into the purge vessel using a 10 μl Hamilton syringe to confirm baseline stability.
14. Record any chemiluminescent peak and allow the signal trace to return fully to baseline before the next injection.
15. Begin standard calibration by injecting 10 μl of the 0.125 μM nitrite solution. Perform two injections for each nitrite standard, recording both peaks that will later be averaged.
16. In the “Comment” tab annotate the sample (for example “10 μl 0.5 μM nitrite”) and press enter. Keep track of the injected samples in this way.
17. After each injection wipe the syringe on a paper towel and rinse 3-4 times with sterile nitrite-free water to remove excess triiodide (The chemiluminescent signals are recorded as peaks in Voltage V over time s).
18. After completing the standard injections, select “Window>Flow Analysis” on the top left of the software. A new screen will appear tabulating the time (s) and height of peaks (V) for each injection.
19. Click and drag the cursor to highlight each peak, adjust to acquire a numerical value of the area under the curve (AUC). After highlighting all the peaks, unclick the “S” next to the tabulated values and select “C”.
20. Change the numbers in the “Amount” column to reflect the known amounts of nitrite in each standard solution.
21. Click “Calibration curve” button at the top left screen (The software will later use this calibration to calculate the amount of nitrite in each tested sample based on the AUC for each sample peak).
22. A window will appear with the calibration curve. If the r^2^ value (the coefficient of correlation between nitrite concentration and area under the signal peak) depicted on the curve is greater than or equal to 0.99, proceed with sample injections.
23. Thaw samples previously collected from ALI cultures on ice.
24. Inject 50 μl of cell-free media, followed by each of basal media sample into the purge vessel and record sample peaks.
25. Wait for 20-30 seconds between injections to let the signal traces descend to the baseline. Do two 50 μl injections per media sample.
26. Annotate each sample in the “Comment” tab with sample ID before injecting. Note: 50 μl injections result in frequent bubble formation, so change the triiodide reagent with anti-foam in the purge vessel regularly.
27. For the apical wash, perform two 50 µl injections per sample and record sample peaks.
28. For the lysate samples perform two injections per sample as done for the basal media and apical wash samples.
29. After finishing the injections, click “Window>Flow Analysis”, and highlight each of the sample peaks as done for the standards.
30. Using the area under the curve (AUC) for the known standard nitrite concentrations, the software calculates the nitrite concentrations for each test sample.

### 7. Data collection, normalization and analysis

1. Export the data obtained through the “Flow Analysis” function in the eDAQ software into an excel file, which tabulates height of each peak (V), area under the curve (V.s) as well as the mean area under the curve for two injections per standard or sample.
2. Also export the raw data of chemiluminescence signal traces from the start to the end of the experiment. This excel file will tabulate the height (V) versus time (s) of signal traces for the entire NOA experiment.
3. Determine the picomoles (pmol) of nitrite injected into the NOA by multiplying the amounts of nitrite (μM) for each sample to the volume of each injection (10 μl for the standards and 50 μl for each of the ALI culture samples).
4. Average the pmol nitrite detected for the appropriate controls (Cell-free media, wash buffer PBS or RIPA buffer) and subtract those from the pmol for the samples detected in each 50 μl injection.
5. Determine the total pmol of nitrite present in each sample collected at the time of cell harvest, by multiplying the pmol of nitrite resulting from NOA injections with the total volume of each sample collected at the start of the experiment; 750 μl for the media, 150 μl for the apical wash and around 100 μl for the lysate (additionally, for lysates multiple the toral nitrite pmol with 2 to account for the dilution factor of 2).
6. Normalize the pmol of nitrite obtained for each sample by dividing these values by the total protein amount (ug) present in the lysate of the corresponding sample, obtained previously through the BCA assay.
7. Average the nitrite levels (pmol/µg) from duplicate injections for each sample.
8. Plot the averaged and normalized nitrite (pmol/µg of total protein) for each sample on the GraphPad Prism software. Perform statistical analysis through students’ t test (one-tailed) to determine any statistical difference in nitrite/NO between experimental conditions (untreated versus drug treated).

## REPRESENTATIVE RESULTS

The overall goal of this procedure is to quantify nitric oxide (NO) levels in differentiated mouse tracheal epithelial cell (MTEC) cultures at air–liquid interface (ALI). Primary cells are isolated and differentiated according to established protocols^**25-26**^. Over the course of 21 days at ALI, the cells are differentiated into a pseudostratified airway epithelium containing ciliated cells^**25-26**^. Levels of ciliation were assessed by immunolabeling for Foxj1 or acetylated α-tubulin. As differentiation progressed, a gradual increase in FoxJ1 expression within the nuclei of differentiating cells was detected (Fig. 1A), accompanied by a corresponding increase in the number of cells carrying multiple cilia labelled by acetylated α-tubulin (Fig. 1B). By ALI day 21, approximately 10% of the airway epithelial cells display motile cilia (Fig. 1), consistent with previous reports^**25-26**^.

**Figure 1.**
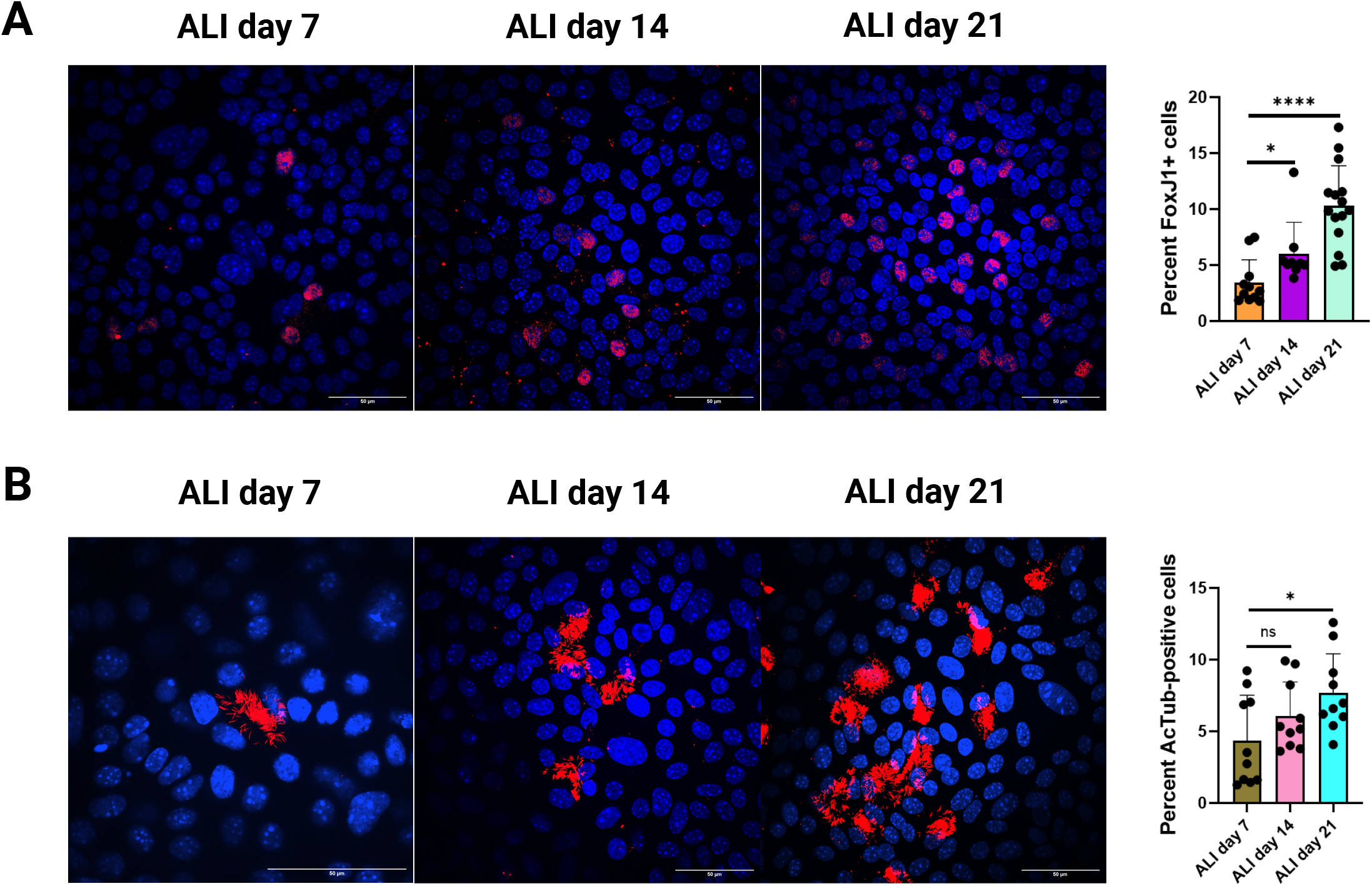
Ciliated cell population increases progressively with ALI differentiation. Mouse tracheal epithelial cells (MTECs) differentiated at ALI at various time points and immunostained with anti-FoxJ1 antibody (A) or anti-acetylated Tubulin (B). Nuclei were counterstained with DAPI. Images are 3D confocal projections. Representative images from 10-15 different regions of interest (ROI) are shown. Scale bar = 50 µm. On the right, quantification of the percentage of cells expressing Foxj1 or acetylated tubulin at different states of ALI. Each bar is Mean ± SD (n=10-15 technical repeats from a single culture). Statistical analysis was performed using Student’s t test (one-tailed); ns, not significant; ^*^ p < 0.05 and ^****^; p < 0.0001.

We previously described the use of ozone-based chemiluminescence for detection of NO in biological samples^**7,16,19-20**^. Similarly, in this study we applied this method for the quantification of nitrite, the primary stable oxidation product of NO, as an indicator of the NO present in samples at the time of collection. In this assay, samples collected from airway epithelial cell cultures are injected into a purge vessel containing the triiodide reagent, where the nitrite (NO_2_^-^) present in the sample is protonated to nitrous acid and subsequently reduced by iodide to yield stoichiometric amounts of NO (Eq. 1). The released NO is then carried by an inert gas, such as helium or argon, into a chemiluminescent detector, where it reacts with ozone to produce excited nitrogen dioxide (Eq. 2). As this molecule returns to ground state, it emits a photon that is detected by a photomultiplier tube, amplified, and converted into voltage signal^**21-22**^ (Fig. 2A).

**Figure 2.**
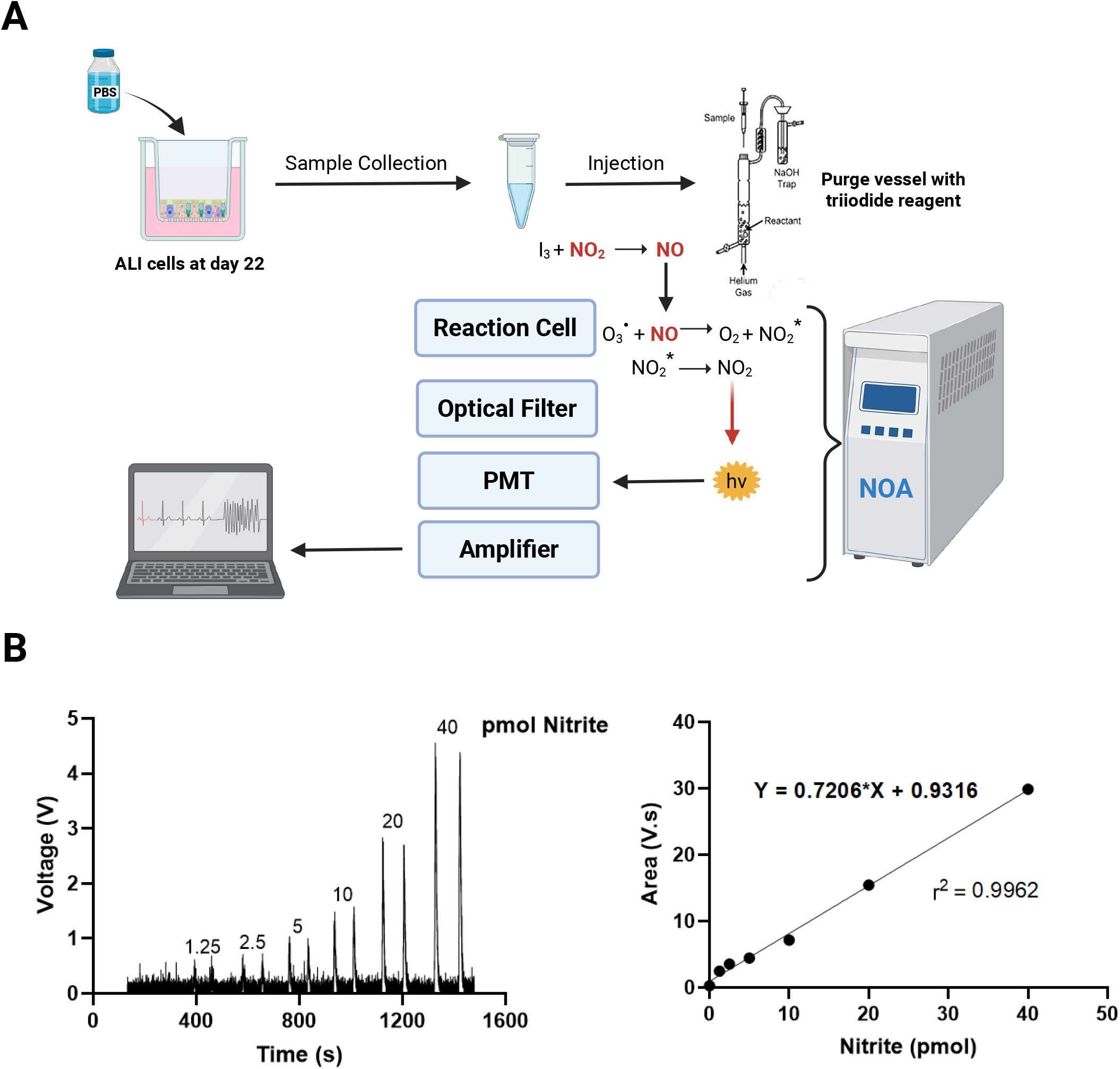
NOA chemiluminescent signals increase linearly with nitrite concentration. (A) Schematic depicting triiodide based chemiluminescent detection of nitrite/NO using the nitric oxide analyzer (NOA) in samples (basal media, apical wash and cellular lysates) collected from ALI-differentiated airway epithelia. Image created with BioRender.com. B) Trace peaks of nitrite standards with each injection at 10 uL volume into the Purge vessel (left). Calibration curve (right) depicts the mean area under the curve for each nitrite standard solution correlating with the picomoles (pmol) of nitrite injected into the NOA.

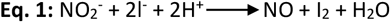

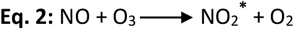

To generate a calibration curve, standard sodium nitrite solutions prepared in PBS are analyzed under identical conditions (Fig. 2B, left). Upon injection into triiodide, the reaction proceeds immediately, producing a voltage response proportional to the total NO derived from nitrite in the standard solution. Chemiluminescent signal traces increase linearly with rising nitrite concentrations. A valid calibration curve, expressed as mean area under the curve (AUC) versus picomoles (pmol) of nitrite, should yield a correlation coefficient (r^2^) greater than 0.99, as shown in Fig. 2B (right).

This procedure was used to analyze samples from airway epithelial cells. As proof of concept that we are indeed measuring NO (via nitrite), we treated cells with N(ω)-Nitro-L-arginine methyl ester (L-NAME), the specific inhibitor of NO synthases (NOSs)^**27**^, and with the NO donor Diethylenetriamine NONOate (DETA/NO)^**17**^. Airway epithelial cells were harvested at ALI day 22 for NO measurements. We first collected samples from basal media and apical washes containing apical mucus secreted by the cells. After removing a small aliquot of the lysate for protein quantification, the remaining lysate was also analyzed. Following calibration, samples were injected into the NOA; each sample was injected twice. The pmol nitrite was averaged for two injections per sample. The averaged pmol nitrite/NO were normalized by protein content in the corresponding cell lysate to obtain nitrite levels (pmol/ug) from at least three independent experiments and values were plotted in GraphPad Prism software (Fig. 3),

**Figure 3.**
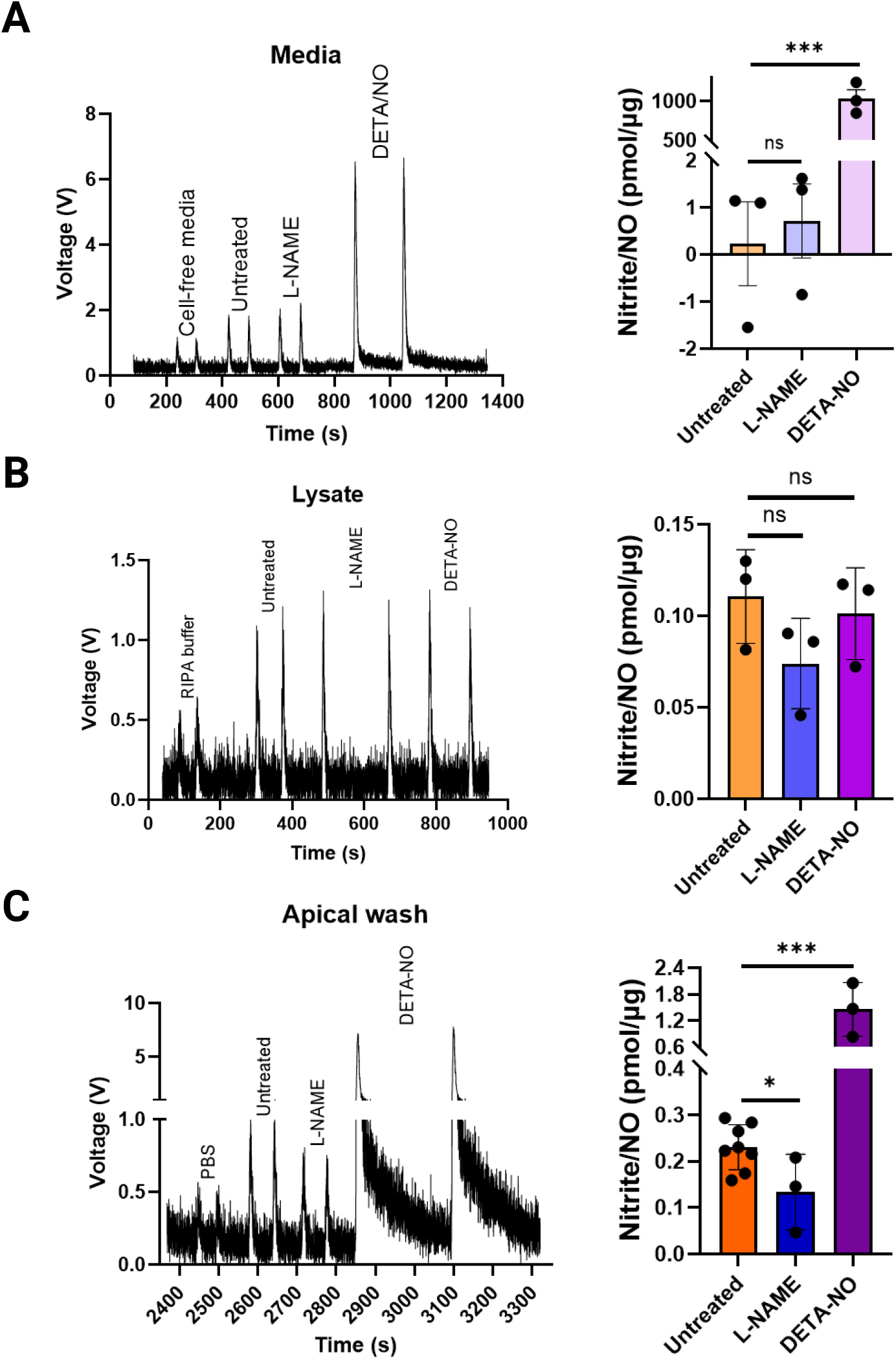
Triiodide based chemiluminescent assay detects nitrite/NO in samples from ALI differentiated airway epithelia. (A) Trace peaks (left) and quantification (right) of nitrite/NO detected in the basal medium of ALI-differentiated airway epithelium. (B) Trace peaks (left) and quantification (right) of nitrite/NO in the cellular lysate. (C) Trace peaks (left) and quantification (right) of nitrite/NO secreted and trapped in the overlying mucus collected as apical washes. Each sample was injected twice with each injection at 50 µl. The detected picomoles (pmol) of nitrite/NO were normalized to the total μg of protein and averaged for duplicate injections. Each bar represents Mean ± SD nitrite/NO levels normalized to pmol/μg protein (n = 3-8, cultures from independent experiments). Statistical analysis was performed using Student’s t test (one-tailed); ns, not significant; ^*^ p < 0.05; ^***^ p < 0.001.

Our results show that in the basal medium, the amount of nitrite/NO is non-significantly altered upon L-NAME treatment compared to untreated controls. In contrast, the positive control DETA/NO treatment shows an expected increase due to NO release from the compound over the 24-hour incubation period (Fig. 3A). In the cell lysate the amount of nitrite/NO was slightly decreased in the presence of L-NAME although not statistically significant (0.112 ± 0.02 pmol/ug in untreated versus 0.074 ± 0.02 pmol/ug L-NAME-treated cells) while in the presence of DETA/NO (0.101 ± 0.025 pmol/ug DETA/NO) levels remained unchanged (Fig. 3B).

Importantly, we observed a significant decrease in nitrite/NO levels in the apical washes from L-NAME treated cells (0.231 ± 0.07 pmol/ug in untreated versus 0.134 ± 0.09 pmol/ug in L-NAME treated) (Fig. 3C). Additionally, DETA/NO treatment resulted in more than 6-fold increased (1.457 ± 0.62 pmol/ug), demonstrating robust nitrite/NO detection on the apical side. This polarized pattern of NO distribution likely reflects the physiology of ALI-differentiated epithelia, where NO production and diffusion are directed toward the apical compartment where mucus efficiently traps nitrite as a stable end-product. The minimal change in basal medium or lysate fractions may be due to intracellular scavenging, dilution, or limited NO retention in these compartments.

Overall our data confirms that the mucus secreted by airway epithelial cells and collected in the apical wash is a reliable reservoir reflecting NO release from ciliated cells, validating the sensitivity and accuracy of the current chemiluminescent method. By enabling direct quantification of NO in an in vitro system that accurately recapitulates airway physiology, this approach offers unique potential to dissect NO-related mechanisms and therapeutic modulation in airway disease.

## DISCUSSION

Multi-ciliated cells within the airway epithelium are critical for NO synthesis and signaling, which in turn is essential for regulating ciliary beating, mucociliary clearance, and airway homeostasis^**6-8**^. NO is a highly reactive molecule and even small changes in NO levels are associated with airway diseases^**1-4**^. Therefore, maintaining finely tuned NO levels is essential for normal airway function. In this study, we describe a method for detecting NO produced by ciliated airway epithelial cells that can be used as an in vitro model of airway diseases associated with dysregulated NO signaling. Mouse tracheal epithelial cells (MTEC) were isolated and differentiated at ALI, with slight modifications from previously published methods^**25-26**^, to obtain a sufficient population of multi-ciliated cells needed to produce measurable nitrite/NO levels (Fig. 1A, B).

Nitrite in biological samples originates primarily from the oxidation of NO by oxygen or reactive oxygen/nitrogen species such as superoxide and peroxynitrite^**19-20**^. Minor contributions can arise from enzymatic pathways involving xanthine oxidoreductase, myeloperoxidase, or hemoglobin/myoglobin, which can oxidize NO to nitrate or reduce nitrite back to NO under certain redox conditions^**28**^. Although nitrite can also be generated by nitrate reduction or by decomposition of reactive nitrogen species, in the presence of normal oxygen levels, these reactions contribute minimally under physiological conditions to total nitrite amount and the vast majority of nitrite in cells derives from NO oxidation^**18,29**^. For future mechanistic studies, it will be important to investigate the potential involvement of other nitrogen species that may influence ciliary function. However, for the purpose of this protocol, which is focused on monitoring NO levels, quantifying nitrite offers a reliable and representative measure, as most of it originates from NO.

Because NO levels in these samples are expected to be very low, relatively large injection volumes are required to achieve detectable signal intensity. The low yield of secreted NO/nitrite from each 12-mm transwell insert necessitates pooling the apical washes from three inserts per condition to reach detection levels in the NOA. However, sample volume must be carefully controlled, as excessive injection of biological material, particularly from lysates with high protein content, can cause protein-induced bubbling in the purge vessel, leading to overflow into the NOA gas lines. To prevent this, the injection volume is maintained at approximately 50 µL into 4 mL of triiodide solution, which provides a balance between signal sensitivity and instrument safety. ALI differentiation for a longer period (28-35 days, for example) can possibly increase the secreted NO in each insert due to a higher number of multi-ciliated cells. Alternatively, bigger commercially available inserts (24 mm diameter) that can seed a greater number of airway epithelial cells can be used to address the issue of low nitrite/NO yield per insert. It is worth noting from a method standpoint that all the apical wash samples collected from pooling three inserts were injected into the NOA in duplicate. Importantly, we analyzed biological replicates, and the results shown here represent data from at least three different independent cultures.

The main concern when employing a highly sensitive method to measure nitrite/NO is the possibility that the detected nitrite originates from external contamination rather than endogenous NO production. Because nitrite is ubiquitous in the environment, contamination can be difficult to avoid. Therefore, it is essential to run appropriate controls alongside experimental samples. To control for nitrite contamination in the media, we incubated cell-free media in the basal chambers of empty (cell-free) inserts and used these as blanks to subtract background signal in the NOA readings. Similarly, PBS and RIPA buffer samples prepared at the time of apical washes and cell lysis were collected and analyzed in parallel. Particularly in the basal medium, we detected an elevated nitrite background likely originating from serum components and protein additives, which can either contain trace nitrite/nitrate or undergo slow spontaneous oxidation of amino acids and peptides that generate nitrite over time. This explains why nitrite levels in cell-free basal medium can be as high as untreated samples. Consequently, this buffer is not ideal for NOA measurements, as the high background can mask small biological differences.

In addition to controlling for exogenous nitrite, it is important to include both positive and negative experimental controls. We inhibited NO synthesis employing the NOS inhibitor L-NAME to reduce NO signals^**27**^ and applied the NO donor DETA/NO to release NO^**17**^. When all reagents are freshly prepared and samples immediately frozen prior to NOA analysis, these controls effectively minimize the likelihood of false readings due to contamination. The only samples showing statistically significant differences consistent with treatment expectations were the apical washes. This finding reflects the polarized nature of airway epithelia, where NO generated by the cells predominantly diffuses toward the apical compartment and becomes oxidized to nitrite, which accumulates in the mucus over time. In the basal medium no clear differences were detected, also due to substantial endogenous nitrite contamination. In the cell lysate the measured nitrite/NO in the DETA/NO treated cells were unexpectedly low, suggesting that most of the released NO is rapidly directed apically and trapped as nitrite rather than retained intracellularly. Thus, the apical wash represents the most reliable compartment for assessing NO release and validating the sensitivity of this detection method. Together with ongoing efforts to elucidate signaling pathways regulating cilia formation and motility, this approach offers a valuable tool to define the role of NO in airway function and disease, and to guide therapeutic strategies aimed at modulating NO bioavailability and signaling.

## ACKNOWLEDGEMENTS

We thank Dr. Mark T. Gladwin (Dean and Professor, University of Maryland School of Medicine) for guidance on study design. We additionally thank the University of Maryland School of Medicine’s Confocal Microscopy Core– Baltimore, Maryland for use of their Nikon W1 spinning disk confocal microscope (S10 OD026698). This work was supported by National Institutes of Health (NIH) grant 5R01HL168775 to Dr. Paola Corti.

## Table of Materials

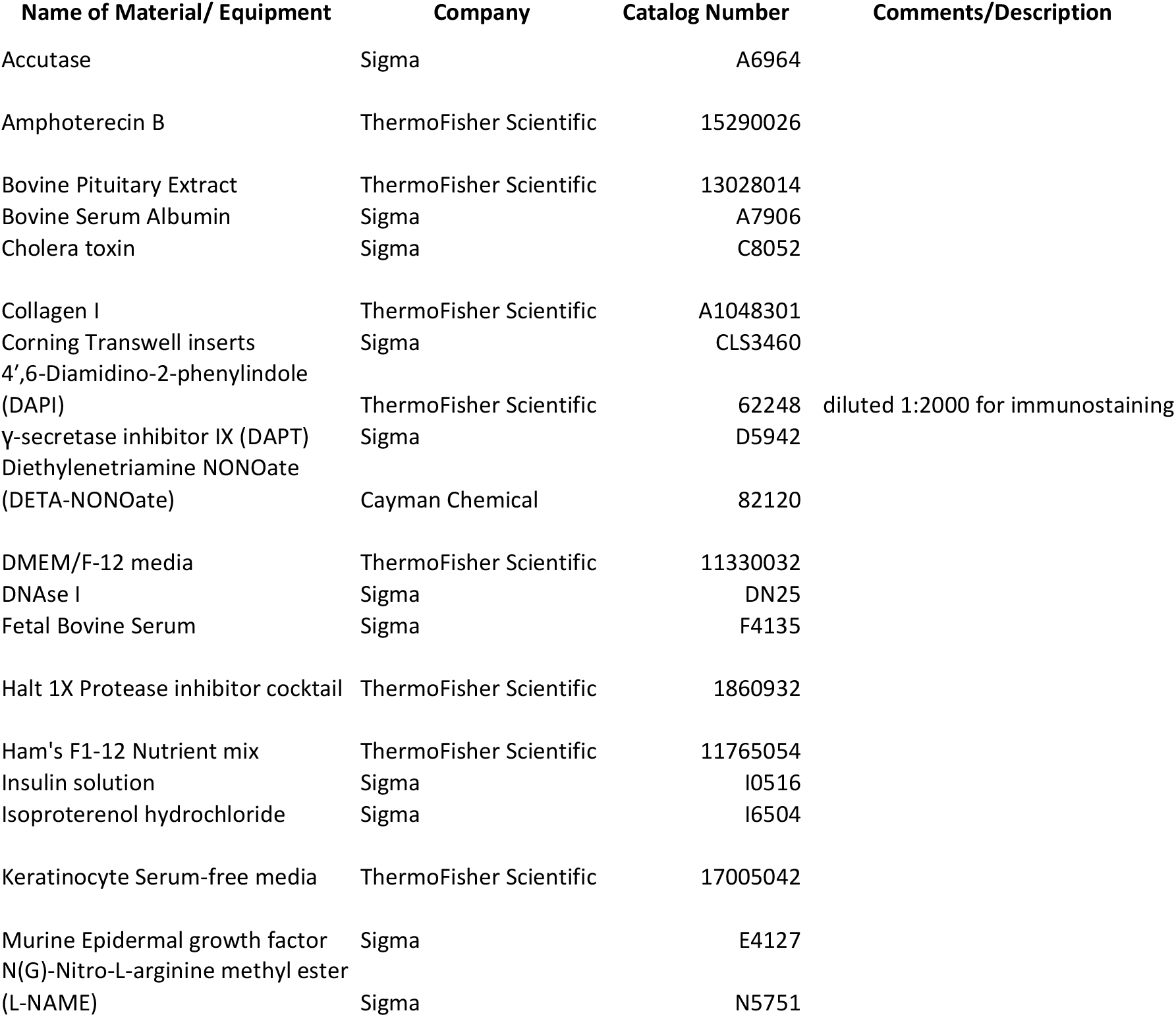
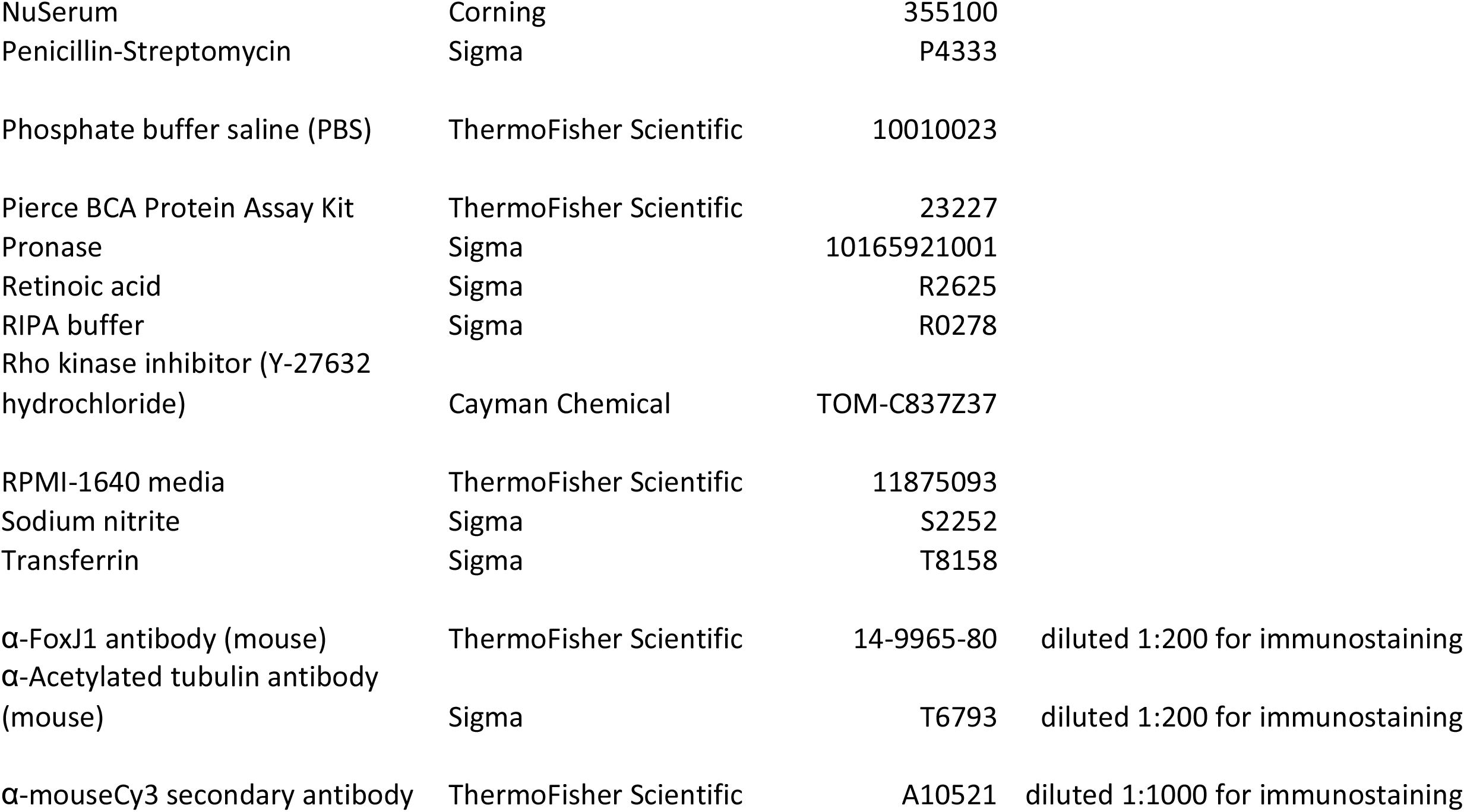

## REFERENCES

1. Alcázar-Navarrete, B., Ruiz Rodríguez, O., Conde Baena, P. Romero Palacios, P. J., Agusti, A. Persistently elevated exhaled nitric oxide fraction is associated with increased risk of exacerbation in COPD. Eur Respir J 51(1): 1701457 (2018).

2. Krantz, C., Janson, C., Alving, K., Malinovschi, A. Nasal nitric oxide in relation to asthma characteristics in a longitudinal asthma cohort study. Nitric Oxide 106:1–8 (2021).

3. Thomas, S. R., Kharitonov, S. A., Scott, S. F., Hodson, M. E., Barnes, P. J. Nasal and exhaled nitric oxide is reduced in adult patients with cystic fibrosis and does not correlate with cystic fibrosis genotype. Chest 117(4):1085–9 (2000).

4. Shapiro, A. J. et al. Nasal Nitric Oxide Measurement in Primary Ciliary Dyskinesia. A Technical Paper on Standardized Testing Protocols. Ann Am Thorac Soc 17(2): e1–e12 (2020).

5. Legendre, M., Zaragosi, L. E., Mitchison, H. M. Motile cilia and airway disease. Semin Cell Dev Biol 110:19–33 (2021).

6. Li, D., Shirakami, G., Zhan, X., Johns, R. A. Regulation of ciliary beat frequency by the nitric oxide-cyclic guanosine monophosphate signaling pathway in rat airway epithelial cells. Am J Respir Cell Mol Biol 23(2): 175–81 (2000).

7. Rochon, E. R. et al. Cytoglobin regulates NO-dependent cilia motility and organ laterality during development. Nat Commun 14(1): 8333 (2023).

8. Mikhailik, A. et al. nNOS regulates ciliated cell polarity, ciliary beat frequency, and directional flow in mouse trachea. Life Sci Alliance 4(5): e202000981 (2021).

9. Schamberger, A. C., Staab-Weijnitz, C. A., Mise-Racek, N., Eickelberg, O. Cigarette smoke alters primary human bronchial epithelial cell differentiation at the air-liquid interface. Sci Rep 5: 8163 (2015).

10. Gindele, J. A. et al. Intermittent exposure to whole cigarette smoke alters the differentiation of primary small airway epithelial cells in the air-liquid interface culture. Sci Rep 10(1): 6257 (2020).

11. Ghosh, B. et al. Effect of sub-chronic exposure to cigarette smoke, electronic cigarette and waterpipe on human lung epithelial barrier function. BMC Pulm Med 20(1): 216 (2020).

12. Baldassi, D., Gabold, B., Merkel, O. Air-liquid interface cultures of the healthy and diseased human respiratory tract: promises, challenges and future directions. Adv Nanobiomed Res 1(6): 2000111 (2021).

13. Bessa, M. J. et al. In Vitro Toxicity of Industrially Relevant Engineered Nanoparticles in Human Alveolar Epithelial Cells: Air-Liquid Interface versus Submerged Cultures. Nanomaterials (Basel) 11(12): 3225. (2021).

14. Allouche, Y. et al. Comparison of submerged and air liquid interface exposure: limitations and differences in the toxicological effects evaluated in bronchial epithelial cells. Environ Res 279(Pt 2): 121856 (2025).

15. Li, J., LoBue, A., Heuser, S. K., Leo, F., Cortese-Krott, M. M. Using diaminofluoresceins (DAFs) in nitric oxide research. Nitric Oxide 115:44–54 (2021).

16. Corti, P. et al. Globin X is a six-coordinate globin that reduces nitrite to nitric oxide in fish red blood cells. Proc Natl Acad Sci U S A 113(30):8538–43 (2016).

17. Zhao, X. J. et al. Mechanisms for cellular NO oxidation and nitrite formation in lung epithelial cells. Free Radic Biol Med 61:428–37 (2013).

18. Lundberg, J. O. Weitzberg E, Gladwin MT. The nitrate-nitrite-nitric oxide pathway in physiology and therapeutics. Nat Rev Drug Discov 7(2):156–67 (2008).

19. Rochon, E. R., Corti, P. Globins and nitric oxide homeostasis in fish embryonic development. Mar Genomics 49:100721 (2020).

20. Giordano, D., Verde, C., Corti, P. Nitric Oxide Production and Regulation in the Teleost Cardiovascular System. Antioxidants (Basel) 11(5):957 (2022).

21. MacArthur, P. H., Shiva, S., Gladwin, M. T. Measurement of circulating nitrite and S-nitrosothiols by reductive chemiluminescence. J Chromatogr B Analyt Technol Biomed Life Sci 851(1–2):93–105 (2007).

22. Basu, S., Ricart, K., Gladwin, M. T., Patel, R. P, Kim-Shapiro, D. B. Tri-iodide and vanadium chloride based chemiluminescent methods for quantification of nitrogen oxides. Nitric Oxide 121: 11–19 (2022).

23. Brody, S. L., Yan, X. H., Wuerffel, M. K., Song, S. K., Shapiro, S. D. Ciliogenesis and left-right axis defects in forkhead factor HFH-4-null mice. Am J Respir Cell Mol Biol 23(1): 45–51 (2000).

24. Jain, R. et al. Temporal relationship between primary and motile ciliogenesis in airway epithelial cells. Am J Respir Cell Mol Biol 43(6): 731–9 (2010).

25. Eenjes, E. et al. A novel method for expansion and differentiation of mouse tracheal epithelial cells in culture. Sci Rep 8(1): 7349 (2018).

26. You, Y., Richer, E. J., Huang, T., Brody, S. L. Growth and differentiation of mouse tracheal epithelial cells: selection of a proliferative population. Am J Physiol Lung Cell Mol Physiol 283(6): L1315–21 (2002).

27. Pfeiffer, S., Leopold, E., Schmidt, K., Brunner, F., Mayer, B. Inhibition of nitric oxide synthesis by NG-nitro-L-arginine methyl ester (L-NAME): requirement for bioactivation to the free acid, NG-nitro-L-arginine. Br J Pharmacol 118(6):1433–40 (1996).

28. Rochon, E. R., Corti, P., Gladwin, M. T. Hemoglobin α in Pulmonary Endothelium: Ironing Out Nitric Oxide Signaling. Am J Respir Cell Mol Biol. 2017 Dec;57(6):639–641.

29. van Faassen, E. E. et al. Nitrite as regulator of hypoxic signaling in mammalian physiology. Med Res Rev. 2009 Sep;29(5):683–741.

